# Soluble FAS Ligand directs human memory B cells towards antibody-secreting cells

**DOI:** 10.1101/2020.12.04.411959

**Authors:** Saskia D. van Asten, Peter-Paul Unger, Casper Marsman, Sophie Bliss, Tineke Jorritsma, Nicole M. Thielens, S. Marieke van Ham, Robbert M. Spaapen

**Affiliations:** Department of Immunopathology, Sanquin Research, Amsterdam, 1066 CX, The Netherlands; Landsteiner Laboratory, Amsterdam UMC, University of Amsterdam, Amsterdam, 1066 CX, The Netherlands; Univ. Grenoble Alpes, CEA, CNRS, IBS, F-38000, Grenoble, France; Swammerdam Institute for Life Sciences, University of Amsterdam, Amsterdam, the Netherlands

**Keywords:** Secretome screen, interferon, FAS, FASL, B cells, germinal center reaction, IgG, differentiation, antibody secretion

## Abstract

Differentiation of antigen-specific B cells into class-switched, high affinity antibody-secreting cells provides protection against invading pathogens but is undesired when antibodies target self-tissues in autoimmunity, beneficial non-self blood transfusion products or therapeutic proteins. Essential T cell factors have been uncovered that regulate T cell-dependent B cell differentiation. We performed a screen using a secreted protein library to identify novel factors that promote this process and may be used to combat undesired antibody formation. We tested the differentiating capacity of 756 secreted proteins on human naive or memory B cell differentiation in a setting with suboptimal T cell help *in vitro* (suboptimal CD40L and IL-21). High-throughput flow cytometry screening and validation revealed that type I IFNs and soluble FAS ligand (sFASL) induce plasmablast differentiation in memory B cells. Furthermore, sFASL induces robust secretion of IgG1 and IgG4 antibodies, indicative of functional plasma cell differentiation. Our data suggest a mechanistic connection between elevated sFASL levels and the induction of autoreactive antibodies, providing a potential therapeutic target in autoimmunity. Indeed, the modulators identified in this secretome screen are associated with systemic lupus erythematosus and may also be relevant in other autoimmune diseases and allergy.

## Introduction

The immune response against a wide variety of pathogens is critically dependent on antigen-specific, high affinity antibodies generated upon natural infection or through vaccination. In contrast, antibodies are detrimental when induced against self-antigens in autoimmune disorders or against allergens, therapeutic proteins, blood products or transplants. Protective and pathogenic antibodies are produced by antibody-secreting plasmablasts or plasma cells (ASCs) that originate from B cells. T cell help consisting of CD40L/ CD40 costimulation and the secretion of specific cytokines is essential for the generation of long-lived plasma cells that produce high-affinity, class switched antibodies. Short-lived plasmablasts differentiated from B cells without T cell help mostly secrete low-affinity antibodies.

T cell-dependent B cells differentiate in secondary lymphoid organs after being activated by their cognate antigen ^1^. Upon activation they migrate to the border of the B and T cell zones where they present antigen-derived peptides to activated T helper cells. After receiving T cell help, B cells migrate back into the B cell follicles while undergoing class switching. They initiate so called germinal center (GC) reactions by alternating between stages of proliferation in GC dark zones (DZ) and reacquisition of antigen to receive additional help from GC-resident antigen-specific follicular T helper (T_fh_) cells in GC light zones (LZ) ^2,3^. Expression of the chemokine receptor CXCR4 allows migration into the DZ, whereas absence of CXCR4 favors LZ localization. During this cycling process, somatic hypermutation of the B cell receptor (BCR) occurs followed by affinity maturation, where B cells with the highest affinity BCRs for the antigen selectively proliferate. Ultimately, these B cells differentiate into memory B cells (CD38^-^CD27^+^) and later into ASCs consisting of plasmablasts (CD38^+^CD27^+^CD138^-^) and terminally differentiated plasma cells (CD38^+^CD27^+^CD138^+^) ^1,4–7^. A subset of memory B cells expresses CXCR3 to migrate into inflamed tissue, while others remain in lymphoid organs or circulates in the peripheral blood ^8^. Upon reinfection and antigen recall, memory B cells may reengage in GC reactions ^8,9^. Plasma cell migration to the bone marrow for long-term survival is directed by CXCR4 ^10^.

Essential for driving B cell differentiation are membrane-bound interactions between receptor-ligand pairs on B cells and T_fh_, such as CD40/CD40L ^11–13^. Furthermore, it is clear that soluble factors like T_fh_ cytokines IL-21 and IL-4 are key T_fh_ cytokines for effective B cell differentiation and that IFN-γ promotes, among other things, migration to inflamed tissue ^11,14,15^. Yet, several other secreted proteins such as IL-10 and type I IFNs affect B cell differentiation in humans, indicating that the role of soluble factors to modulate this process is underexplored ^16–18^.

To study B cell differentiation *in vitro*, T cell help may be mimicked using a CD40L-expressing cell line and recombinant IL-21 and IL-4. This system is well-suited to study which additional signals modulate differentiation of naive or memory B cells into ASCs. Here, we used a previously generated secreted protein library to identify soluble B cell differentiation modulators ^19^. We designed the library to contain immune-related and non-immune human proteins including cytokines, growth factors, peptide hormones and enzymes, because factors from non-hematopoietic cells may also modulate B cell differentiation. Using this diverse library, we identified few additional factors affecting naive B cell differentiation but several molecules with a strong impact on memory B cell differentiation. More specifically, type I IFNs, MAp19 (mannan-binding lectin-associated protein of 19 kDa, transcript variant of complement enzyme MASP-2)^20^ and soluble FASL (sFASL) induced plasmablast differentiation in IgG^+^ memory B cells. Moreover, sFASL promoted the secretion of substantial amounts of IgG_1_ and IgG_4_, showing that FASL drives the formation of ASCs. The different modulators identified in our screens improve the understanding of B cell differentiation and may represent new targets for modulation of B cells and antibody production during disease.

## Methods

### Generation of CD40L expressing 3T3 cell line

NIH3T3 fibroblast cells (3T3) cells were cultured in IMDM medium (Lonza, Basel, Switzerland) supplemented with 10% FCS (Bodinco, Alkmaar, The Netherlands), 100 U/ml penicillin (Thermo Fisher Scientific, Waltham, Massachusetts), 100 μg/ml streptomycin (Thermo Fisher Scientific, Waltham, Massachusetts), 2mM L-glutamine (Thermo Fisher Scientific) and 50 μM β-mercaptoethanol (Sigma Aldrich, St. Louis, Missouri). The 3T3 cells were transfected with Fsp I linearized CD40L plasmid (a kind gift from G. Freeman^21,22^) and Pvu I linearized pcDNA3-Neomycin plasmid) using Lipofectamine 2000 Reagent (Thermo Fisher Scientific) according to manufacturer’s protocol. Three days after transfection, the 3T3 culture medium was supplemented with 500 µg/ml G418 (Thermo Fisher Scientific) to select successfully transfected cells. The 3T3 cells were FACS sorted four times for expression of CD40L using an anti-CD40L antibody (clone TRAP1, BD Bioscience, San Jose, California). This resulted in a stable CD40L-expressing cell line that was cultivated in G418 containing selection media to maintain expression. The same batch of 3T3 CD40L-expressing cells was used for all experiments.

### Secreted protein library

The arrayed secreted protein library was generated previously ^19^. Additional conditioned media were generated using the exact same protocol. In brief, HEK293T cells were individually transfected with plasmids encoding for secreted proteins (OriGene Technologies, Rockville, Maryland and GE Health Care, Chicago, Illinois) using polyethylenimine (Polysciences, Warrington, Pennsylvania). Six hours after transfection, medium was replaced with fresh medium (IMDM supplemented with 10% FCS, 100 U/ml penicillin and 100 µg/ml streptomycin). Three days after transfection, conditioned media were collected and stored in ready-to-screen 96 well plates at −80°C.

### Isolation of human B cells

Buffy coats were obtained from healthy volunteers upon written informed consent in conformity with the protocol of the local institutional review board, the Medical Ethics Committee of Sanquin Blood Supply (Amsterdam, The Netherlands). PBMCs were isolated by density gradient centrifugation using Lymphoprep (Axis-Shield PoC AS, Oslo, Norway). CD19+ cells were isolated from PBMCs using CD19 Pan B Dynabeads and DETACHaBEAD (Thermo Fisher Scientific) according to manufacturer’s protocol with purity >99% and cryopreserved.

### In vitro differentiation culture of human naive and IgG memory B cells

One day ahead of B cell culture, CD40L^+^ 3T3 cells were irradiated with 30 Gy and plated at a density of 10,000 cells/well in a 96 flat-bottom plate (Nunc, Roskilde, Denmark) in B cell culture medium, which is RPMI without phenol-red (Thermo Fisher Scientific) supplemented with 5% FCS, 100 U/ml penicillin, 100 µg/ml streptomycin, 2mM L-glutamine, 50 µM β-mercapthoethanol and 20 µg/ml human apotransferrin (Sigma Aldrich, St. Louis, Missouri; depleted for human IgG with protein G sepharose). The following day thawed human B cells were stained with CD19-BV510 (clone 5J25C1, BD Biosciences), CD27-PE-Cy7 (clone 0323, Thermo Fisher Scientific) and IgG-DyLight 650 (clone MH16-1, Sanquin Reagents, Amsterdam, The Netherlands; conjugated using DyLight 650 NHS Ester (Thermo Fisher Scientific) according to manufacturer’s protocol). CD19^+^CD27^+^IgG^+^ memory or CD19^+^CD27^-^IgG^-^ naive B cells were isolated using an Aria II sorter (BD Biosciences). The sorted B cells were then added to the irradiated CD40L^+^ 3T3 cells at a density of 500-25000 cells/well, in the presence of 5 or 50 ng/ml IL-21 (Thermo Fisher Scientific) as indicated. Conditioned medium from the secreted protein library was added at a 1:12 dilution. For the screens, four wells of empty vector and two wells of IL-21 conditioned media were added to each of the 14 plates. Purified recombinant IFN-α, soluble FAS ligand (both PeproTech, London, United Kingdom) and MAp19 were added at indicated concentrations ^23^. After 9-10 days, B cells were analyzed by flow cytometry. The culture supernatants were used for ELISA.

### Flow cytometry

B cells were stained with LIVE/DEAD fixable near-IR dye (Thermo Fisher Scientific), CD19-BV510, CD27-PE-Cy7, CD38-V450 (clone HB7, BD Biosciences), CD138-FITC (clone MI15, BD Biosciences), for 30 minutes at 4 °C. After staining, cells were washed twice with and taken up in PBS containing 1% BSA and 0.01% azide to be measured on an LSRII (BD Biosciences). Data was analyzed using FlowJo software (BD Biosciences).

### ELISA

Levels of total IgG1 and IgG4 in culture supernatants were determined by sandwich ELISA. Maxisorp ELISA plates (Nunc) were coated overnight with anti-IgG1 or anti-IgG4 (2 µg/ml clones MH161-1, MH161-1, MH164-4 respectively, Sanquin Reagents) in PBS. Plates were washed five times with 0.02% Tween-20 (Avantor, Radnor Township, Pennsylvania) in PBS (Fresenius Kabi, Bad Homburg, Germany), and incubated with culture supernatants (diluted in high-performance ELISA buffer, Sanquin Reagents). Plates were again washed five times and incubated for one hour with horseradish peroxidase-conjugated mouse-anti-human-IgG (1 µg/ ml, clone MH16-1, Sanquin Reagents). After a final five-times wash, the ELISA was developed with 100 µg/ ml tetramethylbenzidine (Interchim, Montluçon, France) in 0.11 mol/L sodium acetate (pH 5.5) containing 0.003% (v/v) H_2_O_2_ (all from Merck, Darmstadt, Germany). The reaction was stopped with 2M H_2_SO_4_ (Merck). IgG concentrations were determined from the 450 nm minus background 540 nm absorption (Synergy2; BioTek, Winooski, Vermont) in comparison to a serial diluted serum pool standard in each plate.

### Screen analysis

The percentages of CD27^+^CD38^+^ and/or CD138^+^ cells within the CD19^+^ population were determined using FlowJo software. These statistics were normalized per plate by B-score normalization using R as described by Malo *et al* ^24^. This method normalizes for column and row confounding effects by three iterative subtractions of median row and column values (excluding IL-21 controls) from each individual well value. After performing this median polish, the B-score for each well was determined by division of the median absolute deviation (MAD = median {|well – median(plate)|}). The screen was performed twice, using B cells from two different healthy donors. The final B-score per secreted protein per condition was the mean of these two screens. The cut-off for hit selection was set at 3 times standard deviation of all conditions, excluding IL-21 controls.

### Proliferation assay

Sorted B cells were washed twice with 10 ml PBS and resuspended to a concentration of 2×10^7^ cells/ml in PBS. Cells and 40 µM CellTrace Yellow (Thermo Fisher Scientific) were mixed at a 1:1 ratio and incubated 15 minutes at RT in the dark, vortexing the tube every 5 minutes to ensure uniform staining. Cells were washed twice using a 10 times volume of cold culture medium to end labeling. Thereafter, B cells were cultured according to the protocol described above. At day 4 and day 10 of culture B cells were mixed with at least 2000 Cyto-Cal counting beads (Thermo Fisher Scientific) or CountBright Absolute counting beads (Thermo Fisher Scientific) and prepared for flow cytometry analysis as described above. Absolute B cell counts were determined according to the formula: 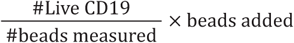.

### Statistics

Statistical analysis was performed using R and GraphPad prism 8.2.1 (GraphPad Software, San Diego, California). *P*-values were determined by indicated statistical tests and depicted using the following symbols: p<0.05 = *, p<0.01 =**, p<0.001 = ***, n.s. = not significant.

## Results

### An in vitro assay allows for unbiased large-scale evaluation of factors required for B cell differentiation into plasmablasts

Because CD40 stimulation and IL-21 signaling seem to be minimal requirements to induce differentiation into antibody secreting cells ^25^, we set out to identify soluble factors that cooperate or act in synergy with these T_fh_ signals. To discover new soluble factors that coregulate human B cell differentiation into antibody-secreting cells, we set up *in vitro* B cell cultures suitable for arrayed secreted protein screening. IgG^+^ memory B cells or naive B cells were cultured with irradiated CD40L-expressing cells in the presence of IL-21 (Fig. 1A) (Unger PP et al., submitted). After nine days, the B cell differentiation state was analyzed by flow cytometry using antibodies against CD27, CD38, CD138 and surface-bound IgG (gating in Fig. S1A). The chemokine receptors CXCR3 and CXCR4 were included in the analysis as they are essential for LZ-DZ cycling and migration to inflamed tissue or bone marrow (Fig. S1A). The B cell differentiation factor IL-21 was titrated to minimize memory and naive B cell differentiation into CD27^+^CD38^+^ ASCs and IgG^+^ B cells respectively, while still supporting B cell survival (Fig. 1B and C). Comparison of the effects of different B cell starting numbers on the efficacy of differentiation into IgG^+^ B cells and CD27^+^CD38^+^ ASCs demonstrated similar trajectories of the IL-21 titration curves (Fig. 1B and C, right panels). Culture initiation with 500 memory B cells per well yielded too few cells for reliable endpoint measurements. Therefore, we chose to start the ensuing differentiation cultures with 1000 memory B cells per well in presence of 5 ng/ml IL-21. For naive B cells, culture initiation with 10,000 cells and 5 ng/ml of IL-21 provided sufficient survival signal for the nine-day culture period, while inducing only few cells to class switch to IgG (Fig. 1C).

**Figure 1.**
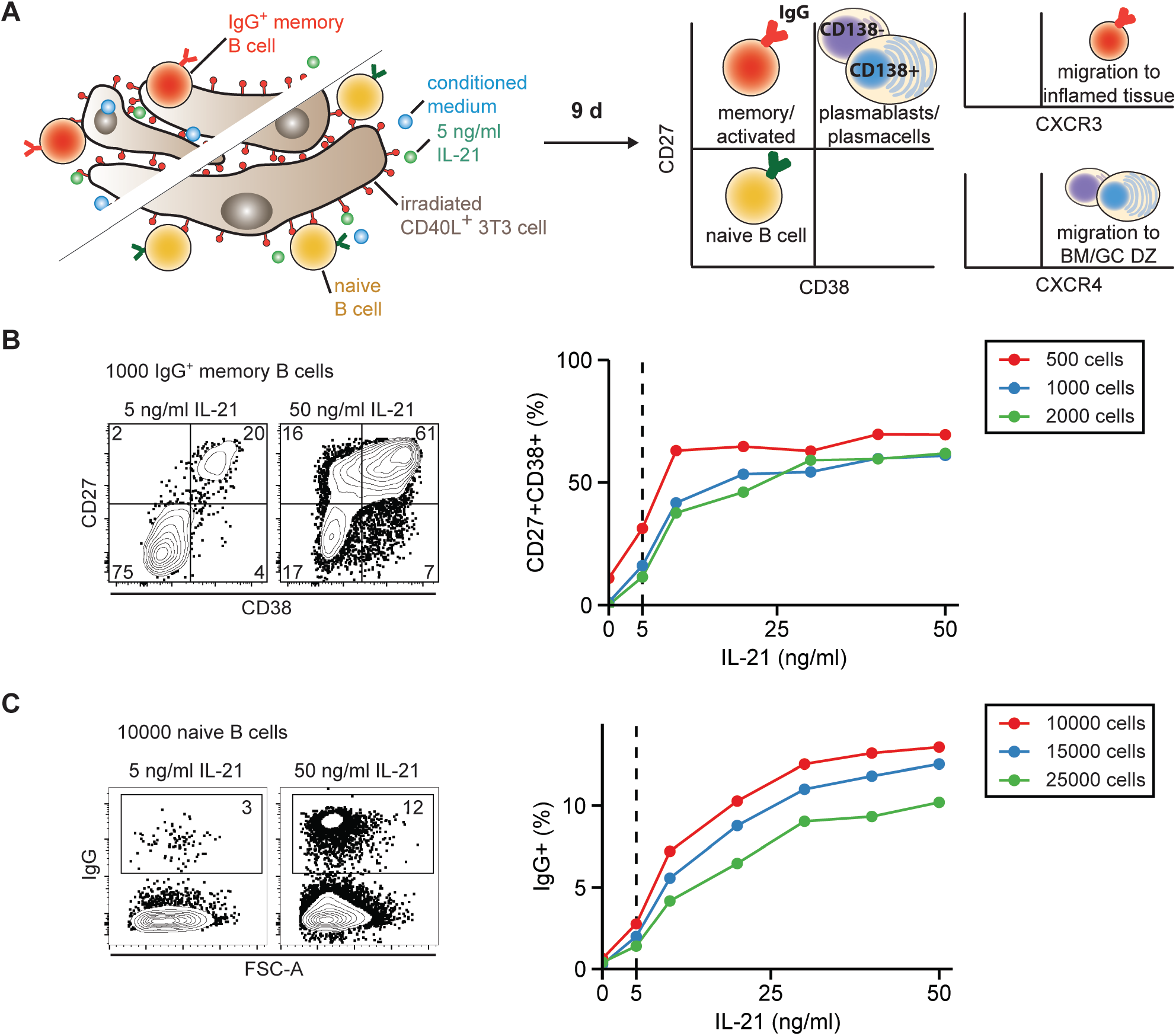
Setup of a suitable assay for arrayed secreted protein screening. (A) Schematic overview of the B cell differentiation assay used in Figures 2-5. Isolated CD27^+^IgG^+^ memory B cells or CD27^-^IgG^-^ naive B cells were cultured for nine days in the presence of irradiated 3T3 cells, 5 ng/ml IL-21 (see B and C) and the individual conditioned media of the secreted protein library. B cells were analyzed by flow cytometry for expression of CD27, CD38, CD138, surface IgG (red), CXCR3 and CXCR4. (B and C) Determination of a suboptimal concentration of IL-21 to culture different indicated starting amounts of memory (B) or naive (C) B cells. Left panels contain example flow cytometry plots of CD27/CD38 (B) and IgG (C) at the indicated concentrations. Right panels show quantification of the percentage of positive cells in the depicted gates. Dashed line indicates 5 ng/ml IL-21. BM = bone marrow, GC DZ= germinal center dark zone.

The screen was executed using a library of arrayed conditioned media each enriched for a single soluble protein secreted by cDNA-transfected HEK293T cells ^19^. This library contains 756 secreted proteins with a broad variety of biological functions, such as neuropeptides, hormones, growth factors and cytokines. To determine the optimal dilution of the library to use in the B cell differentiation screen, we generated IL-21 and empty vector conditioned positive and negative control media respectively. Titrations of both control media showed that the IL-21 conditioned medium was more effective at inducing CD27^+^CD38^+^ ASC differentiation from IgG^+^ memory B cells at lower concentrations, possibly because of supra-optimal IL-21 levels in the conditioned medium (Fig. S1B). While the IL-21 concentration in the supernatant was in the mg/ml range as determined by ELISA (data not shown), we previously found that several other conditioned media in the library contained lower (functional) levels of secreted protein ^19^. For naive B cells, the optimal IL-21 conditioned medium dilution to promote IgG class switching was 1:12 (Fig. S1C). For the library screen on naive and memory B cells, a 1:12 dilution was subsequently used, as it induced no measurable background signal with empty vector (EV) and was sufficient to distinguish IL-21 differentiation effects (Fig. S1B, C dashed line). Finally, we analyzed whether this optimized assay was applicable for screening by determining the dynamic range between 48 negative and positive controls. We found that for several parameters (including the CD27^+^CD38^+^ population and CXCR4 expression) Z’ was higher than zero, indicating that the assay was suitable for large scale screening (Fig. S1D, E).

### IFN-γ induces CXCR3 expression in naive and memory B cells

After setting up a suitable assay, the complete secreted protein library was screened for factors that alter chemokine receptor expression or induce differentiation of B cells. Data from two screens using two different healthy donors were averaged after per plate B-score normalization, which accounts for row and column effects within a single plate as well as variation between plates. Most conditioned media resulted in similar phenotypes compared to empty conditioned media with a B score close to zero (Fig. 2). Positive control IL-21 conditioned medium significantly altered surface CXCR3 and CXCR4 expression in memory B cells and CXCR4 in naive B cells, showing that the library IL-21 conditioned medium was biologically active (Fig. 2A, E, G). The threshold for hit selection was set at three times standard deviation of all conditioned media except the IL-21 positive control (Fig. 2). IFN-γ conditioned medium was the strongest CXCR3 inducer in both memory and naive B cell screens (Fig. 2A-D), in line with previous reports ^14^. In memory B cells, CXCR3 expression was also upregulated by IFN-β1, but not by IFN-α2 (Fig. 2A). Both type I IFNs failed to induce CXCR3 expression in naive B cells (Fig. 2C). In addition, sFASL, FGF5, RLN2, IL15RA and MMP24 conditioned media led to slightly increased CXCR3 levels in memory B cells. None of the conditioned media clearly reduced CXCR4 expression in memory B cells compared to EV (Fig. 2E), yet naive B cells exposed to IFNs, C8B, NRG1, EBI3 and SERPINF1 had lower CXCR4 levels compared to EV (Fig. 2G). HPR, CLC and IL-2 led to marginally enhanced CXCR4 upregulation in memory B cells, and sFASL in both memory and naive B cells (Fig. 2E-H).

**Figure 2.**
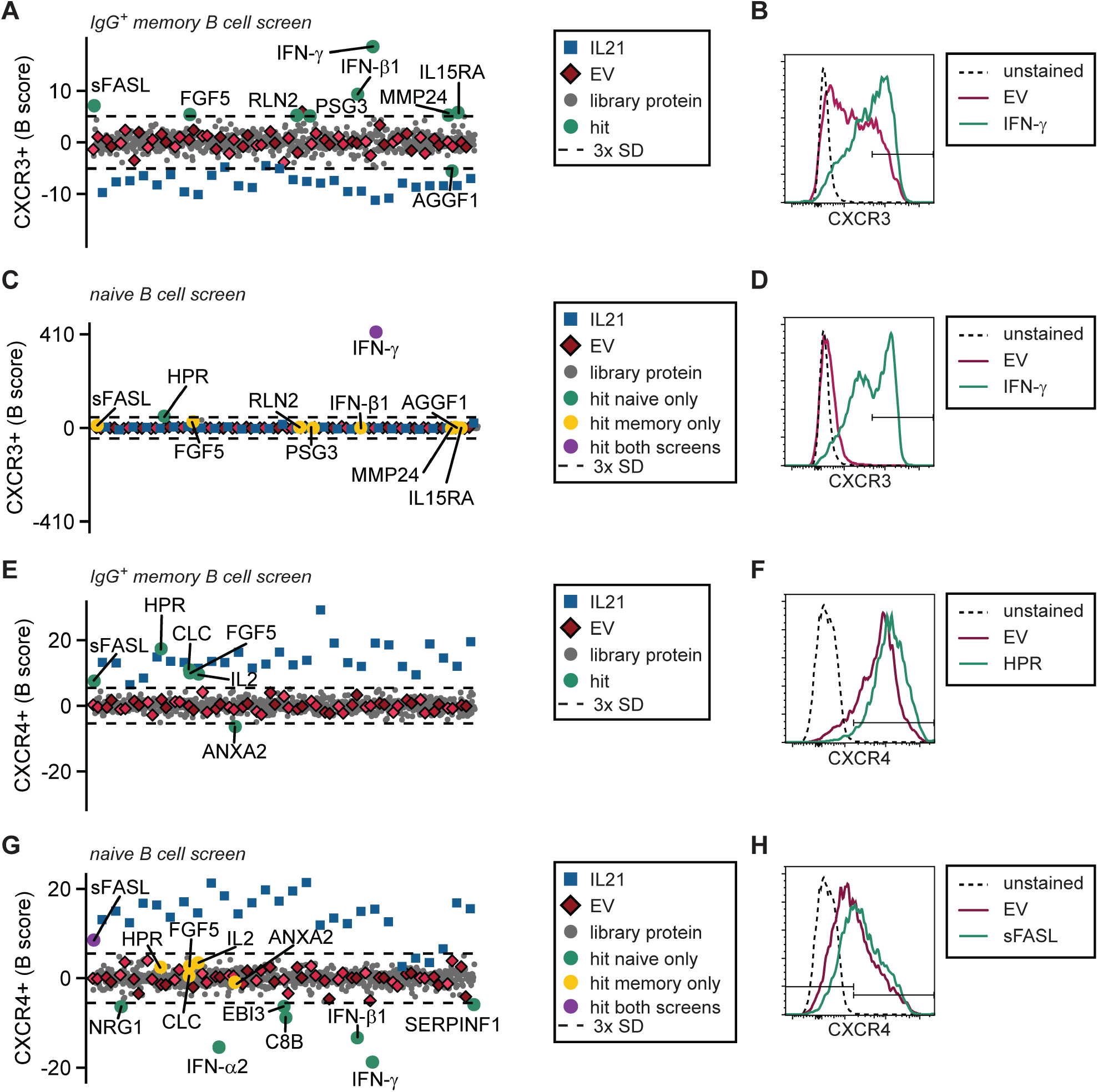
The secreted protein screen identifies several proteins to affect the chemokine receptors CXCR3 and CXCR4. The entire secreted protein library was screened on B cells of two healthy donors. (A-D) Data of CXCR3 expression on IgG^+^ memory (A and B) and naive B cells (C and D). (E-H) Data of CXCR4 expression on IgG^+^ memory (E and F) and naive B cells (G and H). (A, C, E and G) Pooled B score normalized percentages of each phenotype were averaged. For each readout the threshold for hit selection was set at three times SD (dashed line). Controls, hits and other secreted proteins are annotated in the legend boxes. EV is empty vector. (B, D, F and H) Example histograms of the secreted proteins inducing the highest indicated receptor expression on IgG^+^ memory and naive B cells.

### Type I IFNs induce plasmablast differentiation

We next analyzed the capacity of individual secreted proteins to differentiate B cells into CD27^+^CD38^+^ ASCs. Control conditioned medium containing biologically active IL-21 (see Fig. 2) did not greatly induce CD27^+^CD38^+^ ASCs, possibly because of supra-optimal IL-21 levels similar as found during the screen set up (Fig. S1B). Type I IFNs, MAp19 and sFASL all induced memory B cell differentiation into CD27^+^CD38^+^ ASCs, with IFN-α2 and IFN-β1 being the strongest hits (Fig. 3A-C). In addition, naive B cells cultured with type I IFNs differentiated into CD27^-^CD38^+^ B cells (35%), while hardly any CD27^+^ B cells were formed (Fig. 3E-H). The naive B cell differentiation into CD27^+^ B cells was stimulated by IL-2, HPR, CLC, ADAMTS10 and FGF5 conditioned media, but these conditioned media largely failed to drive CD38^+^ plasmablast differentiation.

**Figure 3.**
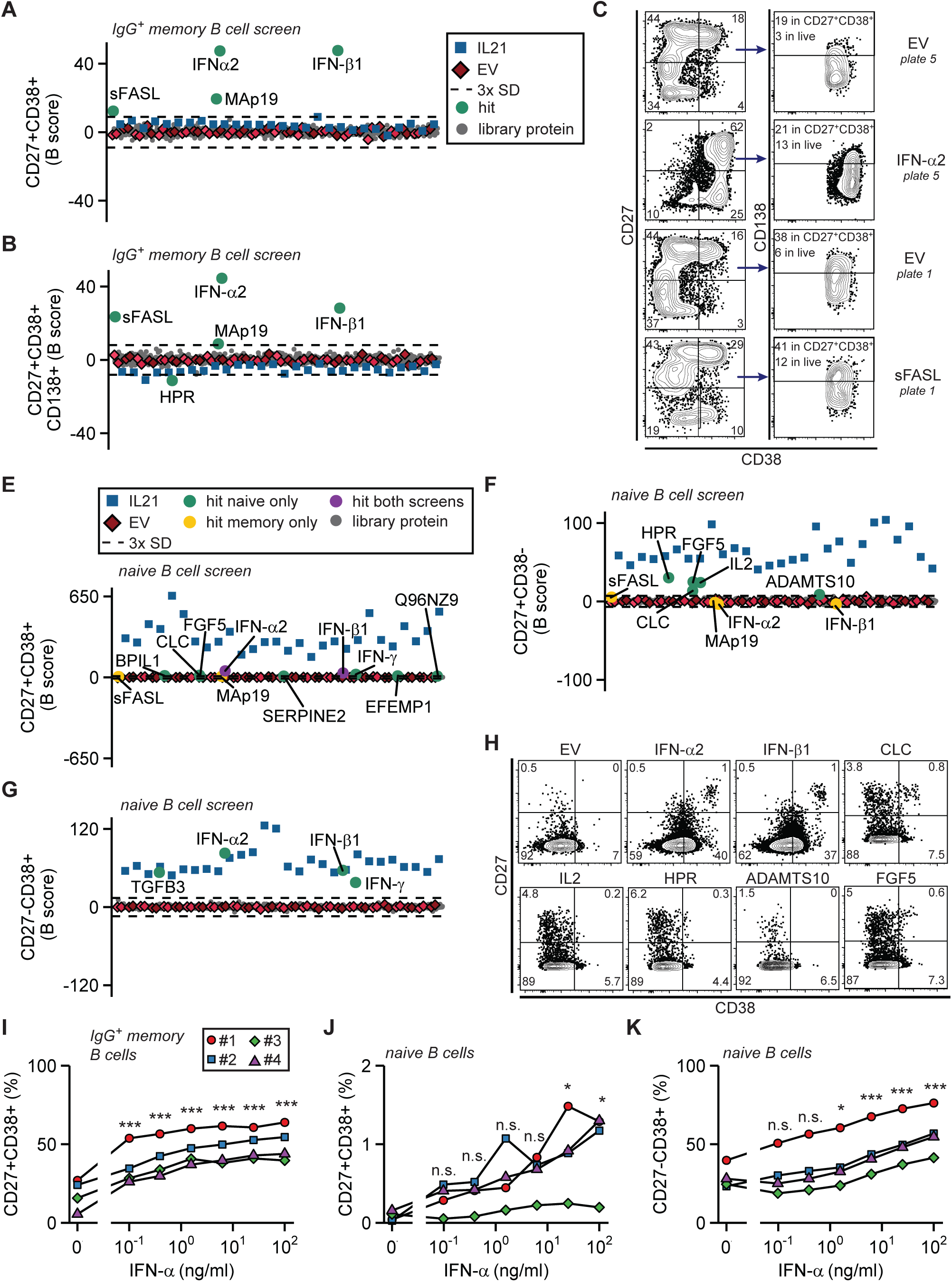
Secretome screening reveals multiple plasmablast differentiation inducing factors. (A, B, E, F and G) Mean B scores of screen readouts for various differentiation stages of IgG^+^ memory and naive B cells from two healthy donors. Legend boxes contain annotation of controls, hits and other secreted proteins. EV is empty vector. Differentiation stage was based on CD27, CD38 and CD138 expression as indicated and gated as shown in the contour plots in (C and H) for controls and the most prominent hits (hit threshold is three times SD; dashed lines). IgG^+^ memory (I) or naive (J and K) B cells from four different healthy donors were cultured as in Figure 1A using serial dilutions of purified recombinant IFN-α instead of conditioned medium. Significance compared to 0 ng/ml IFN-α is shown as determined by ANOVA with Tukey’s post hoc test.

Since type I IFNs were the most potent inducers of B cell differentiation in our screens, we validated these findings using serial dilutions of recombinant purified IFN-α. At all tested concentrations (starting at 0.1 ng/ml), IFN-α significantly directed IgG^+^ memory B cells towards CD27^+^CD38^+^ ASCs (Fig. 3I). IFN-α was also effective in inducing CD38 and CD27 expression in naive B cells, suggesting that the capacity of type I IFN signaling is similar between naive and memory B cells (Fig. 2J-K). Together, our screens reveal that a variety of secreted proteins can stimulate B cell differentiation and support the fact that naive B cells require different signals compared to memory B cells to differentiate into ASCs.

### sFASL induces ASC differentiation

Our screens identified known and unknown secreted proteins to drive naive and memory B cells into various stages of differentiation. As the largest effects were witnessed upon differentiation of memory B cells, we validated these results for two additional donors using newly generated conditioned media. These confirmed that IFN-α2, IFN-β1, MAp19 and sFASL conditioned media induced plasmablast differentiation from memory B cells (Fig. S2). We further continued investigating MAp19 and sFASL, because to our knowledge these factors were unknown to promote B cell differentiation. MAp19 is a splice variant of MASP-2 ^26^. The function of MAp19 is unknown as it lacks the MASP-2 catalytic domain which is responsible for activating complement by cleaving C2 and C4 and may not be sufficiently expressed to compete with MASP-2 for binding to mannan-binding lectin ^20^. FASL is known as an inducer of apoptosis in many (embryonic) developmental and immunological processes ^27,28^. In addition, occasional reports have described cell death independent functions of FASL ^29^. The FASL cDNA in our library encodes for a membrane-bound protein, which may be cleaved to obtain soluble FASL (sFASL) ^30^. As we used the conditioned medium of FASL transfected cells, we hypothesized that sFASL was present in the conditioned media.

Our validation assays showed that the conditioned media containing MAp19 or sFASL significantly induced CD27^+^CD38^+^ B cells in IgG^+^ memory B cells from four different donors (Fig. 4A). To further pinpoint the activity of these proteins, we repeated the assay with purified recombinant MAp19 or sFASL. MAp19 failed to reproduce the conditioned medium effect on B cell differentiation (Fig. 4B). In contrast, sFASL induced B cell differentiation into CD27^+^CD38^+^ B cells starting at 10 ng/ml (Fig. 4C).

**Figure 4.**
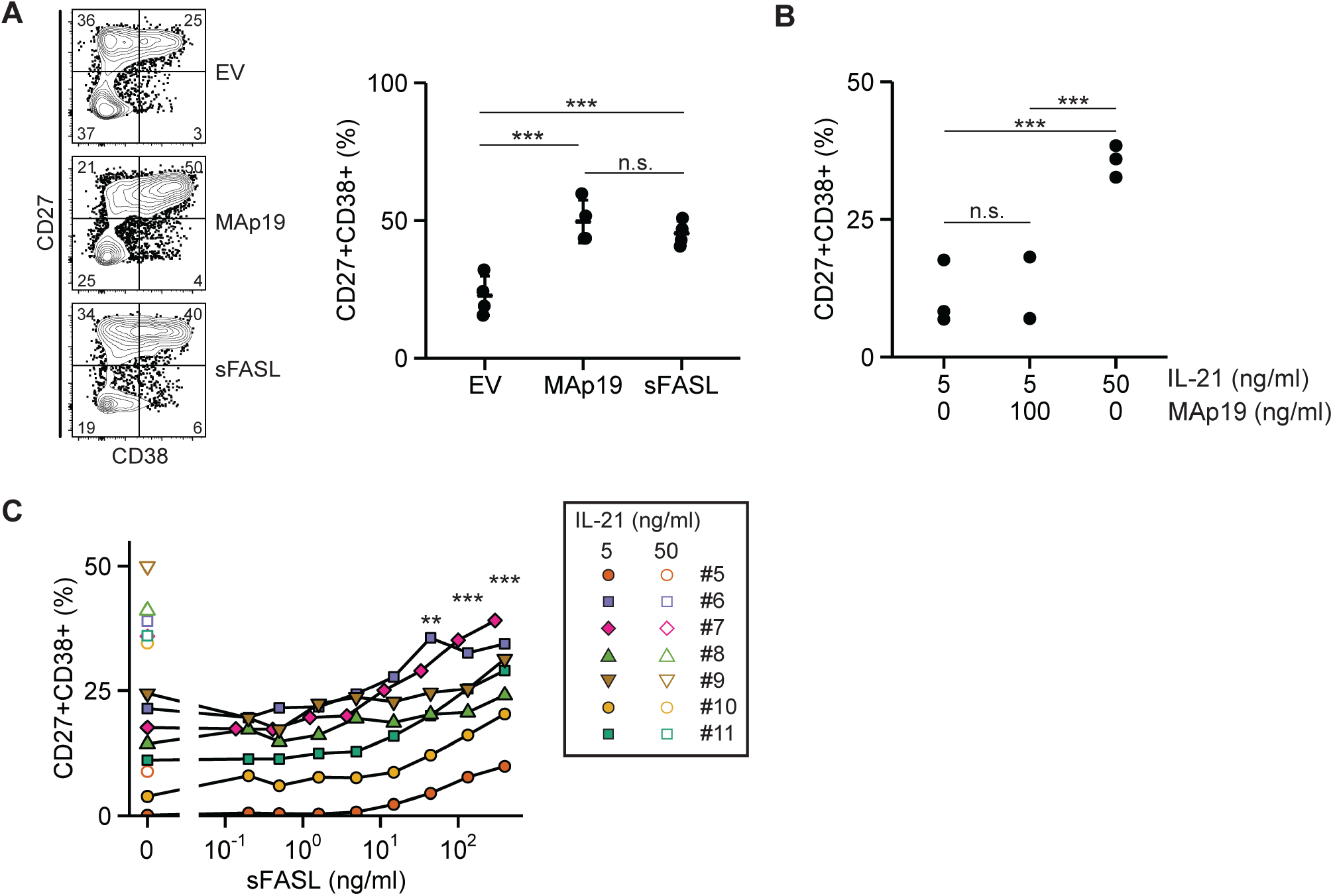
sFASL induces differentiation into ASCs. (A) Newly prepared conditioned media for MAp19 and sFASL were tested for CD27^+^CD38^+^ ASC induction potential on IgG^+^ memory B cells of four different healthy donors (two are the same as in Fig. S2). (B and C) IgG^+^ memory B cell differentiation into CD27^+^CD38^+^ ASCs in the presence of purified recombinant MAp19 (B) or a serial dilution of commercially available recombinant sFASL (C). Statistical significance was determined by ANOVA with Tukey’s post hoc test. In panel only C statistical significance compared to 0 ng/mL FASL is shown.

There are multiple possible explanations for sFASL-driven plasmablast differentiation. sFASL may cause the death of non-differentiating cells, it could encourage the proliferation of differentiating cells or finally, sFASL could drive differentiation into antibody-secreting cells. To gain more insight into the mechanism, we analyzed the effect of sFASL on cell survival and proliferation by flow cytometry 4 days after culture initiation (gating in Fig. S3A). sFASL-treated B cells were still in an early stage of ASC differentiation, and had proliferated at a similar speed as the control without sFASL. Similar B cell numbers were present in sFASL-stimulated cultures compared to the control on both day 4 and day 10, indicating that sFASL does not promote plasmablast differentiation by affecting B cell apoptosis or proliferation (Fig. 5A-C).

**Figure 5.**
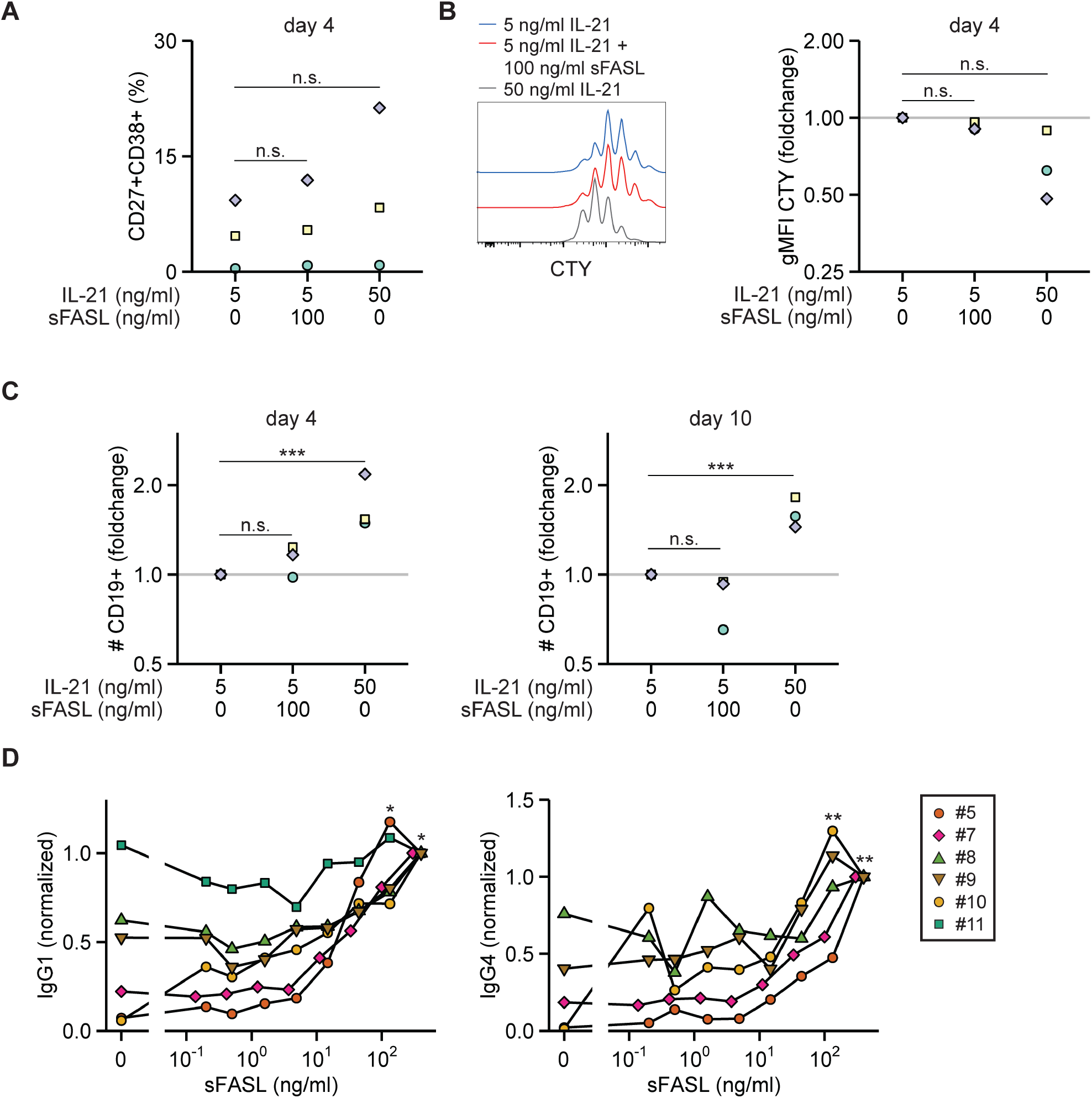
sFASL affects differentiation and not proliferation into IgG1- and IgG4-secreting cells. (A) CD27^+^CD38^+^ expression analysis at day 4 of IgG^+^ memory B cell culture in the presence of purified recombinant sFASL. (B) Histogram of cell trace yellow (CTY) dilution at day 4 after labeling (left panel) and geometric mean fluorescence intensity quantification of CTY dilution from three healthy donors (right panel). (C) Counting bead normalized amount of CD19^+^ B cells after 4-day (left panel) and 10-day (right panel) culture of IgG^+^ memory B cells with indicated conditions. (D) IgG1 and IgG4 levels were determined by ELISA in the culture supernatants obtained at the end of IgG^+^ memory B cell cultures in the presence of serial dilutions of sFASL. Data were normalized to the highest sFASL concentration per donor. Donor numbers (#) correspond to Figure 4C. For donor 11 IgG4 levels were below detection for all conditions including 50 ng/mL IL-21, therefore these data were excluded from analysis. Raw data are plotted in Fig. S3. Statistical significance was determined by ANOVA with Tukey’s post hoc test for panels A-C. In D the Friedman test followed by Dunn’s posttest was applied to determine statistical significance between 0 ng/ml sFASL and the individual serial dilutions of sFASL.

To investigate if these differentiated B cells had become antibody-secreting cells, we measured IgG1 and IgG4 secreted into the culture supernatants. sFASL induced secretion of IgG1 and IgG4 in a concentration-dependent manner (Fig. 5D, S3A-S3B). The levels of produced IgG1 and IgG4 corresponded with the degree of CD27^+^CD38^+^ cell differentiation for each donor (Fig. 4C, S3A-S3B). In conclusion, sFASL induced differentiation of IgG^+^ memory B cells into antibody-secreting CD27^+^CD38^+^ cells.

## Discussion

The germinal center reaction is a dynamic process where activated B cells differentiate into ASCs upon receiving the appropriate signals. By employing a secreted protein library, we successfully identified multiple proteins that promote T cell-dependent differentiation of naive or memory B cells in the presence of to CD40L and IL-21.

The strongest inducers of B cell differentiation in our screens are the well-studied type I IFNs. These multipotent proinflammatory cytokines signal through the Interferon-alpha/beta receptor (IFNAR) to inhibit viral replication by inducing an anti-viral state in non-immune cells and by activating immune cells including B cells. Type I IFNs induce upregulation of CD69, CD86 and CD40 in murine B cells, promote antibody secretion during influenza infection in mice, and can induce antibody secretion in human B cells *in vitro* ^18,31–34^. Although the pro-inflammatory effects of type I IFNs are beneficial during viral infection, type I IFNs may also promote auto-reactive B cells in autoimmune diseases such as systemic lupus erythematosus (SLE) as reviewed by Kiefer *et al* ^35^.

While IFN-α and sFASL directly stimulate differentiation of memory B cells into CD27^+^CD38^+^CD138^+^ plasma cells, purified MAp19 does not. A possible explanation as to why MAp19 conditioned medium induces B cell differentiation and its purified form does not may be that MAp19 stimulates HEK293T cells to secrete another protein or metabolic products. Alternatively, there may be structural differences between the freshly secreted and purified MAp19. Currently MAp19 has no known function ^20^, yet the fact that it is a spliced variant of the complement factor MASP-2 warrants future research on potential regulation of B cell differentiation through the complement system.

FASL is well-known for its apoptosis-inducing capacity. For example, membrane-bound FASL is expressed by activated T cells to kill infected or malignant target cells by binding to and oligomerizing its receptor FAS present on these cells. Apoptosis plays a central role in selection of B cells during affinity maturation. CD40 ligation stimulates FAS expression and sensitizes B cells for FAS-mediated apoptosis, although IL-4 protects B cells from cell death ^37,38^. Yet, in vivo studies showed that FAS signaling is dispensable for GC B cell selection in mice ^39,40^.

The function of sFASL is still under debate as it is less effective in inducing apoptosis than membrane-bound FASL. sFASL may compete with membrane-bound FASL for FAS-binding thereby blocking apoptosis and even promoting proliferation ^30,36^. We did not observe a role of sFASL in the induction of apoptosis. Our results rather highlight a novel function of sFASL, which is to directly induce differentiation of CD40-activated memory B cells into ASCs.

sFASL may partake in the pathogenesis of several autoimmune diseases. Increased sFASL levels have been found in the serum of SLE patients and saliva of Sjögren’s syndrome patients compared to healthy individuals. In addition, patients with more severe rheumatoid arthritis have higher sFASL levels in synovial fluid compared to those with less severe pathology ^41–43^. These autoimmune diseases are all characterized by the presence of autoantibodies, produced by ASCs originating from self-reactive B cells. So far, it remains unclear if the relationship between sFASL levels and autoimmunity is causal. Our data now demonstrates a direct role for sFASL in promoting B cell differentiation. It therefore opens the way for novel research exploring if sFASL initiates or worsens immune-mediated diseases by promoting differentiation of detrimental, autoreactive B cells.

## Acknowledgments

The authors thank the Sanquin Research Facility for their assistance with flow cytometry. This research was supported by Sanquin Product and Process Development of Cellular Products grants PPOC-14-46 to Dr. Robbert Spaapen, and both the PPOC-17-34 and the Landsteiner Foundation for Blood Transfusion Research (LSBR) grant 1609 to Dr. Marieke van Ham. IBS acknowledges integration into the Interdisciplinary Research Institute of Grenoble (IRIG, CEA).

