## Supplementary Figures for "Soluble FAS Ligand directs human memory B cells towards antibody-secreting cells"

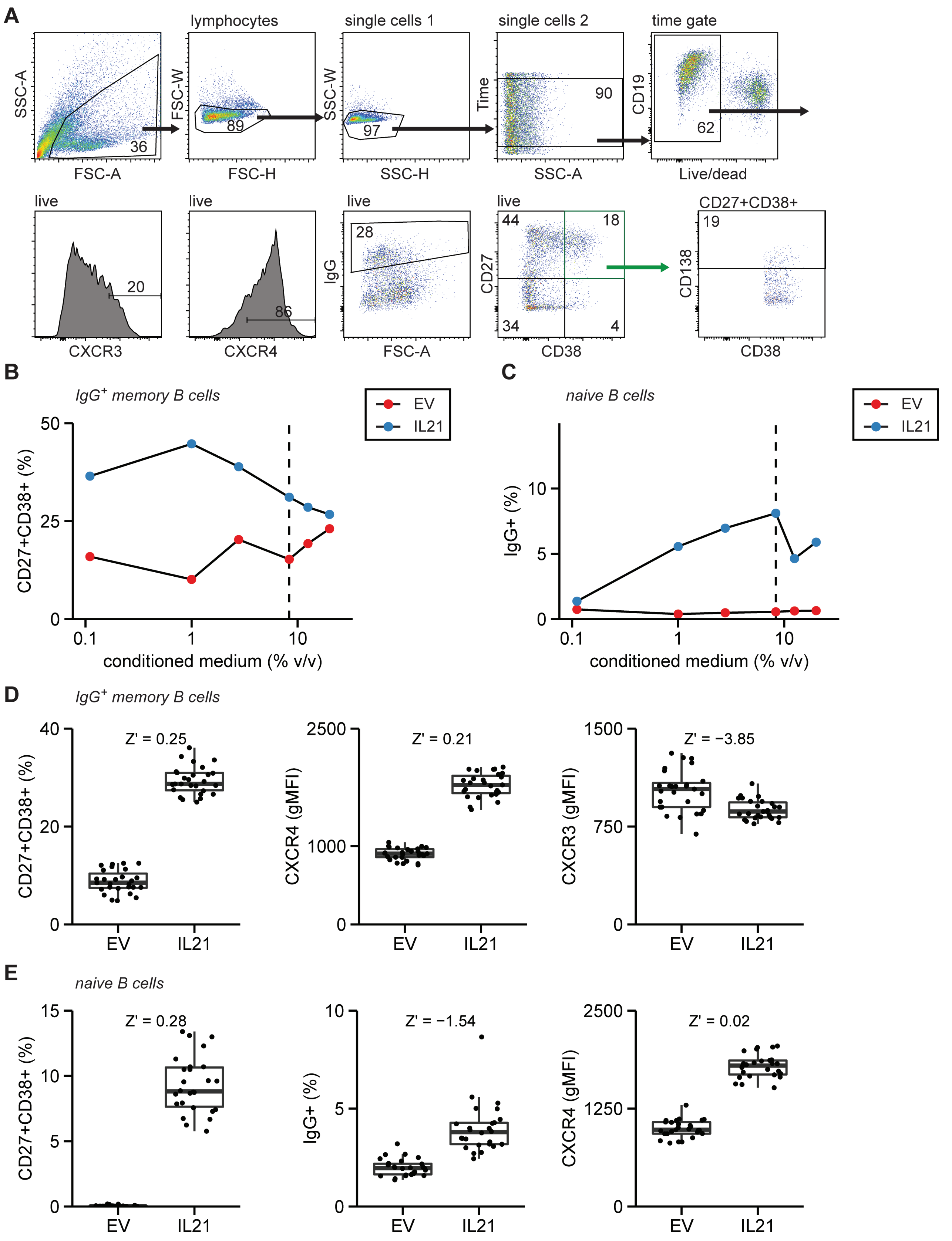


**Supplementary Figure S1. Optimization of B cell differentiation assay for arrayed screening** (A) Strategy at the end of the B cell differentiation assay to gate live, single CD19+ B cells. Markers of interest within these B cells include CXCR3, CXCR4, IgG and also CD27/CD38 to allow gating on plasmablasts and plasma cells. Plasma cells were identified by gating for CD138 within CD27$^+$CD38$^+$ B cells. Percentages of CD27$^+$CD38$^+$CD138$^+$ cells were calculated within live B cells. (B and C) IgG$^+$ memory (B) and naive B cells (C) were cultured as depicted in Figure 1A in the presence of a serial dilution of conditioned medium derived from empty vector (EV) or IL-21 cDNA transfected HEK293T cells in addition to 5 ng/ml IL-21. 1:12 (8.3 % v/v) was chosen as the preferred conditioned medium dilution for the secreted protein screen, here indicated with a dashed line. (D-E) 24 replicates of control empty vector and IL-21 conditioned media were assayed according to the optimized conditions. Depicted parameters were readout at the end of IgG$^+$ memory (D) and naive B cell (E) cultures. Z' scores are shown, Z'>0 indicates suitability of the assay for arrayed screening.

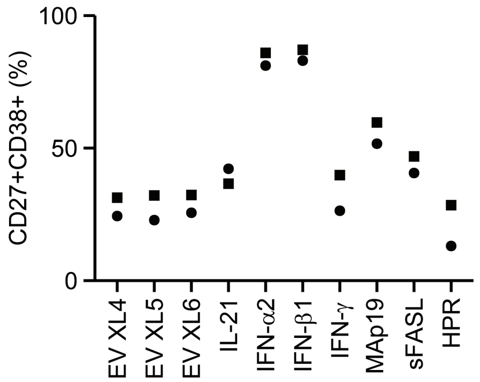


**Supplementary Figure S2. Type I IFNs, sFASL and MAp19 induce the formation of CD27⁺CD38⁺ ASCs.** Newly prepared conditioned media from indicated cDNAs and empty vectors (XL4, XL5 and XL6 backbones) were tested for CD27⁺CD38⁺ ASC induction potential on IgG⁺ memory B cells of two different healthy donors (part of the data is also included in Fig. 4A).

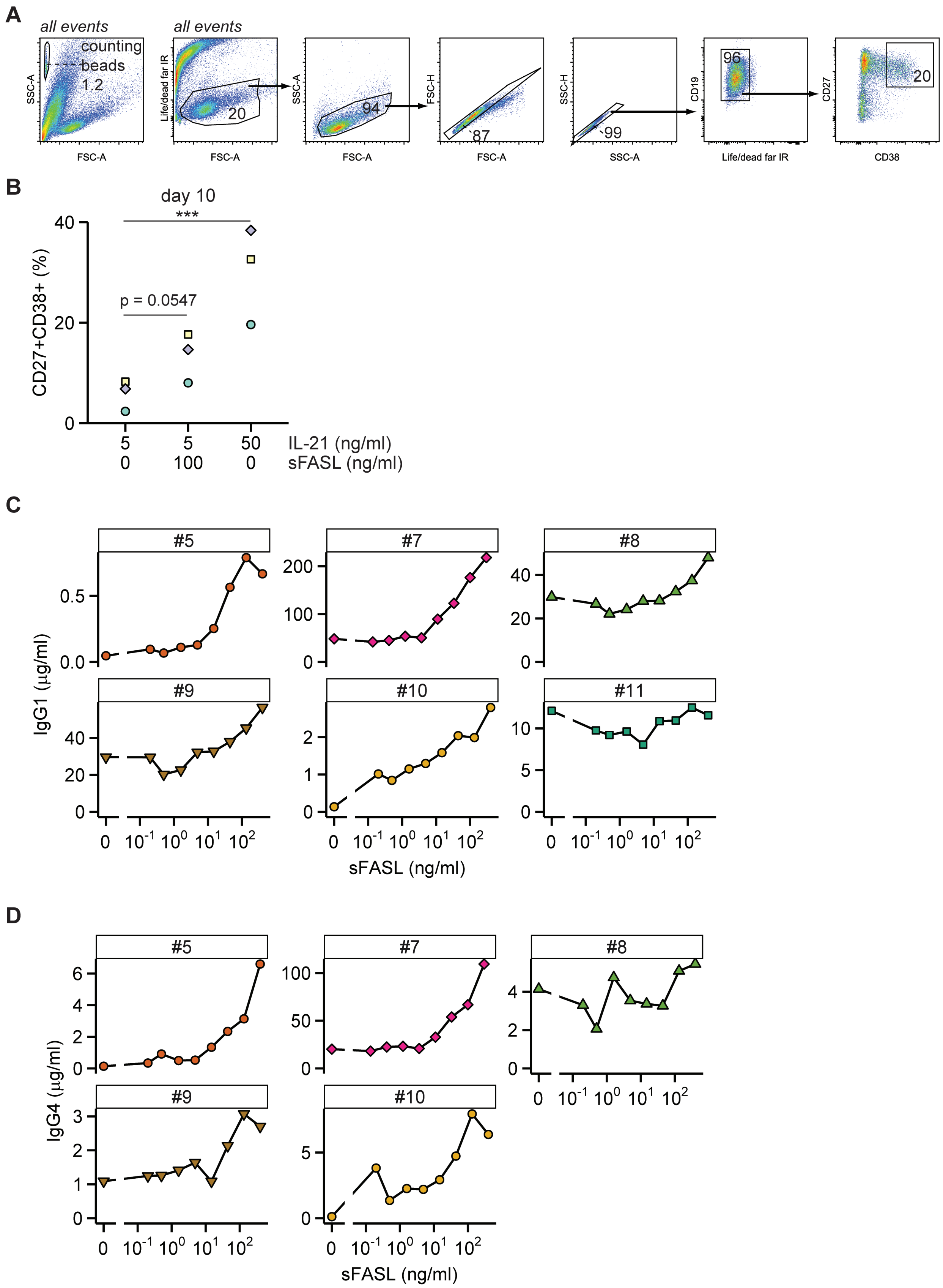


**Supplementary Figure S3. Gating strategy of B cell proliferation assay** (A) Strategy to gate on counting beads, $CD19^{+}$ live cells and $CD27^{+}CD38^{+}$ ASCs on both day 4 and day 10 of the proliferation assay. (B) Quantification of $CD27^{+}CD38^{+}$ ASCs within live $CD19^{+}$ B cells on day 10. Statistical significance was determined by ANOVA with Tukey's post hoc test. (C and D) IgG1 (C) and IgG4 (D) levels in culture supernatants of $IgG^{+}$ memory B cells cultured in the presence of sFASL serial dilutions.
